# Insights into mortality patterns and causes of death through a process point of view model

**DOI:** 10.1101/067967

**Authors:** James J. Anderson, Ting Li, David J. Sharrow

## Abstract

Process point of view models of mortality, such as the Strehler-Mildvan and stochastic vitality models, represent death in terms of the loss of survival capacity through challenges and dissipation. Drawing on hallmarks of aging, we link these concepts to candidate biological mechanisms through a framework that defines death as challenges to vitality where distal factors defined the age-evolution of vitality and proximal factors define the probability distribution of challenges. To illustrate the process point of view, we hypothesize that the immune system is a mortality nexus, characterized by two vitality streams: increasing vitality representing immune system development and immunosenescence representing vitality dissipation. Proximal challenges define three mortality partitions: juvenile and adult extrinsic mortalities and intrinsic adult mortality. Model parameters, generated from Swedish mortality data (1751-2010), exhibit biologically meaningful correspondences to economic, health and cause-of-death patterns. The model characterizes the 20^th^ century epidemiological transition mainly as a reduction in extrinsic mortality resulting from a shift from high magnitude disease challenges on individuals at all vitality levels to low magnitude stress challenges on low vitality individuals. Of secondary importance, intrinsic mortality was described by a gradual reduction in the rate of loss of vitality presumably resulting from reduction in the rate of immunosenescence. Extensions and limitations of a distal/proximal framework for characterizing more explicit causes of death, e.g. the young adult mortality hump or cancer in old age are discussed.

## Introduction

The vast majority of models that characterize historical patterns of mortality and project future trends are based on age and time dependent changes in the rate of mortality from all causes (Booth and Tickle 2008). Such models do not explicitly address the underlying processes leading up to the mortality event and are therefore not equipped to address the interactions of proximal (i.e. acute) and distal (i.e. chronic) causes of death (COD). Applications of rate of mortality models to project cause-specific mortality assume independent causes and over predict the combined mortality (Alai et al. 2015). Furthermore, projecting future mortality trends based on historical trends of specific COD patterns is problematic when underlying COD changes over history (Tuljapurkar 1998). Finally, mortality rate based approaches are less equipped to incorporate the rapidly expanding biological knowledge on aging and mortality. An alternative modeling approach, which takes the process point of view (POV) (Aalen and Gjessing 2001), by tracking the processes leading up to a mortality event has properties that, in principle, are better equipped to address the proximal and distal interactions of COD using simplified representations of biological processes of aging and mortality^1^.

The theme of this paper is to explore cellular and system level processes that contribute to mortality and expand a vitality model based on the process POV. We first review development of process POV models of mortality and the historical development of vitality, a central tenant of the models. We next review recent studies on cellular and physiological mechanisms of aging and mortality and highlight their vitality-like properties. We then discuss leading COD in this framework, emphasizing the importance of proximal and distal processes in COD classifications. With this background we expand an existing two-process model with the inclusion of a juvenile mortality component and discuss candidate biological mechanisms for the model terms. Finally, we fit the model to the 250-year Swedish mortality database and discuss the historical trends in the model coefficients and mortality components in the context of the effects of historical changes in environmental, social and health factors on the putative biological processes the model represents.

## Process point of view of mortality

Process POV models of mortality are not new to demography and were the basis of the first models published in the 19^th^ century. Gompertz (1825) introduced the first process POV model in which mortality was assumed to be the result of stress challenges exceeding the innate survival capacity or vital power which declines with age. Makeham (1867), in an insightful discussion, laid the foundations for mortality components by partitioning mortality challenges, e.g. diseases, into aggregates: one of sufficient strength to overcome any level of vital power and a second weaker aggregate in which mortality depends on the remaining vital power. Strehler and Mildvan (1960) mathematically captured the process of mortality as challenges to vital power in terms of the probability of a challenge of random magnitude and frequency exceeding a linearly declining survival capacity, denoted vitality. An alternative form of mortality was introduced with first passage time models. In the most common form, based on the Wiener process, the time to mortality is determined by the stochastic passage of vitality into a zero boundary denoting death or failure (Aalen and Gjessing 2001; Anderson 1992; Anderson 2000; Li and Anderson 2009; Steinsaltz and Evans 2004; Weitz and Fraser 2001; Whitmore 1986). The probability distribution of first passage time is defined by the mean and variability in the stochastic rate of loss of vitality normalized by the initial vitality. The two mortality processes were combined by assuming that challenges to vitality do not influence the absorption of vitality paths into the zero boundary (Li and Anderson 2013). This two-process model partition of the mortality into two types: extrinsic mortality resulting from challenges to vitality and intrinsic mortality resulting from absorption of vitality into the zero boundary^2^.

### Distal/proximal factors of mortality

In a two-process POV framework the intrinsic/extrinsic partition of mortality also requires a further classification of mortality into distal and proximal factors. The distal factors shape the trajectories of vitality with age by altering the rate of loss of vitality. The proximal factors determine the immediate mortality event. With these definitions, both intrinsic and extrinsic mortalities have distal factors defined by the rate of loss of vitality while the proximal factors have different mathematical forms: vitality challenge event for extrinsic mortality and vitality absorption event for intrinsic mortality. However, functionally an absorption event can also be viewed as a weak challenge event that only results in mortality in low-vitality individuals. Thus, functionally the proximal factor is partitioned on the strength of the challenge: extrinsic mortality challenges are strong and infrequent while intrinsic mortality challenges are weak and frequent. As we explore further on, the biological mechanisms of the two forms are different. The immediate advantage of a distal/proximal framework is in mathematically linking a cause of death and its associated risk factors. For example, pneumonia as a COD, could be partitioned into a proximal factor, a pathogen exposure, and a distal factor characterized by the level of vitality at the time of exposure, which depends on age and previous exposures to stressors (McCabe et al. 2009; Rudan et al. 2008).

### Evolving understanding of vitality

Process POV models reviewed above and other similar models employ an abstract measure of survival capacity that was first expressed as the power to avoid death (Gompertz 1825) and later as vital power (Makeham 1867). Strehler and Mildvan introduced the term vitality, which subsequently became the most common usage^3^. Previously, there was little discussion of underlying physiological and biochemical mechanisms of vitality. However, some information on its nature was gleaned from studies of interventions that affect lifespan. Strehler (1959) compared relationships of radiation exposure and physiological aging on lifespan and concluded they are unlikely to operate through similar mechanisms. An important finding of the study was that a number of measures of human functional capacities, e.g. basal metabolic rate, plasma flow, declined in a linear manner in adults, comporting with the assumption of linearly declining vitality with age. A number of studies in which a stressor dose alters a survival curve provides additional insights into the nature of vitality. Fitting vitality models to survival curves of fish, insects and worms exposed to different stressors demonstrated that vitality loss rate increased linearly with stressor magnitude (Anderson 2000; Anderson et al. 2008; Gosselin and Anderson 2013). A recent study of the effects of a variety of lifespan interventions on *Caenorhabditis elegans* provides more insights. In the study Stroustrup et al. (2016) manipulated lifespan with genetic (genes for heat shock, hypoxia, insulin pathways) and physiological (diet, thermal and oxidative stress) interventions and found that the resulting survival curves, either increasing or decreasing lifespan, could be superimposed onto a single curve by simply rescaling the time axis with the rescaling related to the intervention magnitude. The authors concluded that the temporal rescaling of survival curves with significantly different interventions indicates that diverse biochemical pathways to mortality may converge through a single stochastic pathway that determines the probability of death from the multiple causes. They suggested that the rate of loss of the survival factor depends on the intervention magnitude. In our terminology and mathematical framework this factor is vitality and its rate of loss is a fundamental determinant of lifespan. Additional studies on human mortality patterns have also demonstrated that changes in survival across years and population groups can be characterized by changes in the rate of loss of vitality (Li and Anderson 2013; Li and Anderson 2015; Li et al. 2013; Sharrow and Anderson 2016a; Sharrow and Anderson 2016b).

In summary, from Gompertz to the recent works a theme emerges that complex pathways leading to death can be described by challenges to an age-declining state, which by convention we denote vitality. This framework fits data well, is mathematically tractable and has a general sense of properties of aging^4^. While, the possibility that highly complex pathways leading to death pass through a single pathway as proposed by Stroustrup et al. (2016) is surprising and compelling, its validity is not established by model fits to data. As noted by Steinsaltz and Evans (2004) and others, biological mechanisms underpinning such patterns need to be demonstrated and linked to the model parameters. The difficulty is that studies linking biological mechanisms to vitality have been essentially correlative. Strehler and Mildvan (1960) used linear declines of physiological functions with age to hypothesize that vitality is an index of available energy capacity. They also noted that the correlation did not prove the hypothesis, but alternative variables were unsuccessful so the energy analogy was the most appropriate at the time. The recent studies mentioned above demonstrating interventions quantitatively change lifespan are evidence to suggest that the rate of vitality loss and rate of aging may be equivalent. Thus, to seek a biological basis of vitality we consider the mechanisms of aging.

Although little was known about mechanisms of aging when Strehler and Mildvan developed their vitality-energy analogy, a half century later more than 300 theories of aging had been proposed (Medvedev 1990; Yin and Chen 2005). Currently, the aging field is in flux because of rapidly expanding research in longevity and new insights in evolutionary theories of aging. For example, a central prediction of the evolutionary theory that senescence is driven by extrinsic mortality has been shown to be wrong (Wensink et al. 2016). Other studies suggest evolution of senescence through mutation accumulation proposed by Hamilton (1966) needs to be revised (Wachter et al. 2013) and diverse theories need to be merged to understand evolution of senescence (Wachter et al. 2014). Similar challenges face cellular-level theories of aging. A dominant theory that aging results from molecular damage by reactive byproducts of metabolism is not supported by experiments. An alternative theory (cell hyperfunction) proposes fast nutrient processing that is beneficial in early growth and reproduction becomes detrimental in later life (Gems and Partridge 2013; Kenyon 2010). Furthermore, it has also been proposed that the root cause of both damage and hyperfunction-like processes is the fact that physiochemical reactions are always imperfect and this imperfection cannot be removed by natural selection (Gladyshev 2016). Although it might seem that aging research is in disarray, major molecular pathways of aging have been enumerated and qualitatively linked to mortality and longevity. However, the divergent views of aging as a complexity of molecular pathways vs. aging as a linear decline in population vitality need to be reconciled to advance process POV models and ultimately bridge the conceptual divide between cell aging and population survival.

## Candidate processes underlying vitality

Thus, our goal here, motivated by the consistent support of a process POV framework of mortality, is to draw on the expanding biological knowledge to identify candidate processes underlying vitality. Our approach is akin to that of Strehler and Mildvan, in posing underlying mechanisms that seem most reasonable with the information available. We select our candidate processes based on their cell and system level functions and how these functions change across a lifespan and relate to historical changes in the environment and society. Our underlying premise is that vitality realistically captures patterns of mortality because the distal and proximal factors of the vitality framework capture the aging patterns of cells and physiological systems. We next briefly review these patterns.

### Cellular hallmarks of aging

At the molecular and cellular levels, nine hallmarks of aging have been identified and placed into three hierarchical groups (López-Otín et al. 2013). The first group involves damage of cellular processes (genomic instability, epigenetic alterations, telomere attrition, and loss of proteostasis). A middle group (deregulation of nutrient sensing, mitochondrial dysfunction, and cellular senescence) are compensatory responses that mitigate acute damage but can also exacerbate chronic damage. The final group are integrative hallmarks involving effects of the other two groups on tissue homeostasis and function. They include the decline of regenerative potential of stem cells with age and chronic inflammation of tissue (stem cell exhaustion and altered intercellular communication). From the perspective of the vitality framework, the properties of the first group of hallmarks are similar to the distal factor of mortality, in that as vitality is a declining measure of survival capacity, this first group of hallmarks develops with age, reducing the capacity of the cells to maintain homeostasis. The properties of the second and third groups are similar to the proximal factor of mortality in that the processes transition to a lower, or nonfunctional state, when the distal processes reach a limit. For example, telomeres, the protective end-caps of chromosomes, are akin to vitality in that they shorten with each cell replication. Cell senescence is akin to vitality boundary passage in that when telomeres reach a critical length the cell enters replicative senescence (Blackburn et al. 2015). In a similar manner autophagy, the process to remove damaged proteins and invasive pathogens in cells, is akin to vitality in that it declines with age by the accumulation of waste material. Analogous to vitality challenges, cells enter apoptosis when the rate of pathogen flux or protein damage exceeds the autophagic capacity (Cuervo et al. 2005; Denton et al. 2015; Han et al. 2015). However, while these hallmarks have a mechanistic link to aging, individually they do not have strong correlations with measures of aging (e.g. Arai et al. 2015; Boonekamp et al. 2013; Glei et al. 2016).

### Differential aging in regulatory networks

The hallmarks of aging develop stochastically in physiological systems causing different forms of aging. For example, physiological system blood biomarkers (e.g. lipids, leucocytes, oxygen transport, liver, vitamins and electrolytes) exhibited different age-related deviations from the norms and different patterns with causes of death. Notably, cardiovascular disease was predicted by several of the above mentioned biomarkers but cancer was generally unassociated with any (Li et al. 2015). Additionally, an index of biological markers of aging was a better predictor of mortality risk than an index of activity markers (Cornman et al. 2016). These studies and others suggest that, leading up to death, the dysregulation of the multiple cellular regulatory networks progresses in semi-independent patterns (Cohen 2016; Riera et al. 2016). The important point for a vitality framework is the possibility of representing the distal factors of COD with separate vitality streams associated with different physiological networks.

### Immune system

At the system scale, a major focus of aging involves the immune system, because it is the network of cells, tissues and organs that protect and repair body damage resulting from internal and external stresses and pathogens (Weiskopf et al. 2009). Note also, immune system function involves all of the primary hallmarks of aging (López-Otín et al. 2013). For example, autophagy is the central process of the immune system for engulfing and removing pathogens (Kuballa et al. 2012; Puleston and Simon 2014), telomere attrition limits the ability of the immune system to generate lymphocytes (Aubert and Lansdorp 2008; Blackburn et al. 2015) and epigenetics remolding of the chromatin contributes to the age-dependent increase of the immune system inflammatory response (Booth and Brunet 2016). Two aging hallmarks, the gradual deterioration of the immune system, known as immunosenescence (Aw et al. 2007), and chronic low-grade inflammations, known as inflammaging (Franceschi et al. 2007; Frasca and Blomberg 2015), are central actors in determining death due to infections (Weiskopf et al. 2009), cancers (Bonafè et al. 2001; Palucka and Coussens 2016; Vasto et al. 2009), heart disease (Epstein and Ross 1999; Legein et al. 2013) and other diseases (Chung et al. 2009).

For identifying candidate processes for vitality, we focus on the cells of the innate and adaptive immune systems and how they change with age. The innate immune system, mainly consisting of monocytes, neutrophils, natural killer and dendritic cells, represents the first line of defense against pathogens. The cells are constantly produced by stem cells in the bone marrow and freely circulate but do not divide. The adaptive immune system consists of B and T cells, which are produced by stem cells in the bone marrow and mature as naïve cells in the bone marrow and thymus respectively. Dendritic cells of the innate immune system present antigens of pathogens to naïve B and T cells, which then clonally divide as effector cells to mount further defense. After an infection is removed a fraction of the effector T cells remain as memory T cells, which are able to quickly respond to future infections (Weiskopf et al. 2009). Both innate and adaptive immune cells engulf and digest the invading pathogen and regulate homeostasis through autophagy (Janeway et al. 2005; Levine and Deretic 2007).

### Aging patterns of the immune system

Patterns of mortality over life have been partitioned into two evolutionary-significant stages: ontogenescence, representing the declining mortality rate from conception to maturation and senescence representing the increasing mortality rate from maturation to old age (Levitis 2011). In both stages, robustness is considered a key factor in determining patterns of mortality. Changes in robustness are closely tied to changes in the immune system, which exhibits distinct patterns from birth through childhood, mature adulthood and old age (Simon et al. 2015). In our vitality framework we equate robustness to vitality and our working hypothesis is that the immune system is the key player that changes over age. Therefore, we focus here on the contributions of the immune system to ontogenescence and senescence separately. This partition then becomes the basis for separate vitality streams for child and adult stages. The details are as follows.

#### Ontogenescence

It is well established that fetal and newborn mammals have limited ability to mount immune responses (Holt and Jones 2000). In particular, the limitation involves low innate system activity at birth, making infants susceptible to bacterial and viral infections (Levy 2007). However, the cell levels change rapidly postpartum. Neutrophils and monocytes dominate the innate system at birth and can exceed adult levels before normalizing within a few days. In the adaptive system, both T and B cells numbers are high at birth, increase further over the first year of life and then decrease towards adult levels in school-age children. However, memory T cells, a measure of the adaptive system’s capacity to respond to infection, are very low at birth, increase steadily to age of 5 years and reach adult levels by age of 15 years (Ygberg and Nilsson 2012). Thus, this pattern is support for a separate vitality process characterizing the effects of immune system development on early life mortality.

#### Immunosenescence

The adult immune system gradually degrades with age through immunosenescence (Gruver et al. 2007; Weiskopf et al. 2009). Degradation is most severe in the adaptive immune system as hallmarked by the decrease in naïve T cells and accumulation of memory T cells. The reduction in naïve T cells is in part driven by reduced bone marrow immune cell output involving the involution of the thymus, where T cells mature. The involution begins at puberty and declines by about 3% per year through middle age, 1% in old age and is barely detectable by age 85 (Palmer 2013).

The increase, or inflation, of memory T cells is associated with infections of cytomegalovirus (CMV), which once established, results in an inflation of memory T cells as the immune system tries to contain, but not remove, the infection (Pawelec et al. 2009). In developed countries, a significant proportion of the population is CMV-negative, but with age, the population that is CMV-positive increases, plateauing at about 85-90% by age 75-80. The prevalence of infection is greater in developing countries where 90% may be infected in youth (Pawelec et al. 2012). CMV infections have been associated with a number of chronic illnesses and markedly increase vascular disease COD in individuals over 65 years of age (Savva et al. 2013) but may have beneficial effects on the immune system in young individuals (Sansoni et al. 2014). Besides contributions of thymus involution and CVM infections to T cell dynamics, immune system degradation also involves interconnected responses of many other processes including epigenetic remodeling of cells, telomere shortening and the diversity of naïve T cell receptors (Ongrádi and Kövesdi 2010).

From the perspective of shaping mortality profiles, the end result of immunosenescence is age-increasing susceptibility to novel infections (Kline and Bowdish 2016). Furthermore, a number of behavioral and environmental factors affect the rate of immunosenescence. A major factor is protein-energy malnutrition in which individuals with insufficient protein in their diet experience atrophy of immune system organs and reduction of immune system leucocytes (Beisel 1996). However, diet restriction that reduces body weight without protein-energy malnutrition can increase immune function (Ritz and Gardner 2006). In general, malnutrition is a common cause of immunodeficiency worldwide (Chandra 1997). Obesity is also a factor in immune system function including reduced number and response of T cells and lymphoid atrophy, which result in higher susceptibility to infectious diseases. Furthermore, evidence indicates that response of the immune system to infection diminishes its capacity to respond to future infections (Schaible and Kaufmann 2007). Finally, exercise has been demonstrated to delay or possibly reverse immunosenescence (Turner 2016).

We hypothesize that immunosenescence, as a generic measure of the ability of the immune system to protect the organism, is a central nexus between the biology of aging and vitality. Viewing the rate of loss of vitality as an index of the rate of immunosenescence highlights important features that need to be considered in extensions of the model. In particular, the rate of change of vitality is population specific and depends on the cumulative experience in terms of past infections and health behavior. These factors can, in principle, be represented as nonlethal challenges that effect vitality loss. In the current form of the model sublethal challenges are subsumed into the mean and variability in the rate of loss of vitality. In other words, in estimating vitality rate parameters we implicitly include the effects of random challenges on the rate of loss of vitality. This suggests a contribution of nonlethal proximal factors to shaping the distal factors of mortality thus offers a quantitative framework in which to represent the interplay between life history and the patterns of morbidity and mortality in populations. However, this linkage is not explored in this paper.

A final note of the importance of the immune system in the vitality framework involves its centrality. The immune system consists of a complex of free cells that reach and service all other systems of the body. It embodies the properties of the unknown state variable that Stroustrup et al. (2016) hypothesize is required to produce temporal scaling of lifespan interventions. That is, all other biochemical pathways that affect lifespan pass through the state and the state itself changes with age and the magnitude of intervention. No other physiological system of the body has these properties, and therefore our working hypothesis is that vitality is an aggregate measure of the immune system function. We do note however that a process POV admits multiple vitality streams and in our model below we represent separate streams for immune system development and senescence.

## Cause of death

Here, we consider how challenge and boundary passage events can be related to COD. In particular, using the historical shift in COD from infectious diseases to degenerative diseases and cancers we illustrate that the partition into intrinsic and extrinsic processes of mortality depends on the age of death and the mix of challenge types.

### Infectious disease

The patterns of infectious diseases (ID) through human history have been linked to ecological processes including the spatial distribution, movement and nutritional status of populations (Dobson and Carper 1996). Reductions in mortality from infectious disease can be largely reduced to two factors: reductions in disease transmission and host susceptibility (Cohen 2000). In the context of the vitality framework we propose these factors are expressed by three processes: disease transmission is characterized by the frequency of extrinsic challenges, the disease virulence is expressed by the challenge magnitude and host susceptibility is expressed by vitality.

Beginning the 20^th^ century, infectious diseases were the leading cause of death worldwide. In the United States tuberculosis, pneumonia and diarrhoeal disease accounted for ~30% of mortalities while at the end of the century they accounted for less than 5%. This decline is first attributed to reductions in both challenge frequency and magnitude. The frequency reduction involved reduced disease transmission through improved sanitation, hygiene and safer food and water. The magnitude reduction is attributed to the introduction of antimicrobial agents in the mid-20^th^ century. However, in developing countries infectious diseases currently account for ~25% of the deaths (Cohen 2000). Finally, the reduction can be attributed to lower host susceptibility involving a lowering in the rate of immunosenescence as discussed above. However, reduction in the impacts of disease challenges have not mitigated the effects of immunosenescence in old age. In people ≥ 65 years old, one third of the deaths worldwide are from bacterial infections including lower respiratory tract, urinary tract and skin/soft tissue infections (Kline and Bowdish 2016). In this age group about 85% of the deaths are attributed to influenza and pneumonia (Mouton et al. 2001).

To a first order, young individuals experience the same disease challenges as adults, but the mortality rate is patterned by their immune system development that results in the mortality rate declining in an exponential-like manner through childhood. Challenges involving complications during neonatal preterm births and intrapartum together account for 70% of deaths in the first 6 days after birth. In the neonatal stage (~2 to 4 weeks) deaths are primarily caused by infections (Heron 2015) and over the first five years of life infectious disease, including acute respiratory infections, diarrhea and malaria, account for about two-thirds of all deaths (World Health Organization 2010). Malnutrition is directly or indirectly responsible for half the deaths per year for children under five and malnutrition also worsens the outcome from other infections including tuberculosis, HIV/AIDS and malaria (Schaible and Kaufmann 2007).

#### Disease Latency

In the vitality framework, mortality from infectious disease is attributed to random challenges that occur over short periods of time. However, in reality a latency exists between the challenge event and the mortality. The latency can be on the order of weeks for diseases such as typhoid and pneumonia and months to years for tuberculosis (Control and Prevention 2016). The long tuberculosis latency is a result of its etiology. It is estimated that one-third of the world population is infected with *Mycobacterium tuberculosis* (Flynn and Chan 2001). Those with active infections develop active tuberculosis within 1 to 3 years while a third become clinically latent and are not infectious. In ~10% of the latent group the infection can reactivate. Treatment is largely successful but in developing countries approximately 7% of those under treatment die within a few months (Birlie et al. 2015; Field et al. 2014). Tuberculosis illustrates a further challenge in partitioning mortalities because of its protracted development it has been classified as an underlying cause of death as well as an associated cause of death with other causes being ascribed as the underlying cause (Santo et al. 2003).

In principle, the latency can decrease with lower vitality. However, in the model framework the classification of extrinsic mortality does not depend on disease latency. Instead, it depends on the distribution of mortalities with age. A pattern of exponentially increasing mortality with age contributes to the intensity of extrinsic mortality in the model.

### Injury

Between ~10–25 years of age, the rate of mortality increases mainly from physical injury associated with risky behavior. Known as the early adult mortality hump, globally in 2004 it accounted for an 82% increase of the total mortalities in this age range (Patton et al. 2009). The phenomena involves a mismatch in the rate of development of the neural systems that control reward seeking and risk avoidance behaviors (Shulman et al. 2016). In the vitality framework this increase can be expressed as a transient increase in challenge frequency and magnitude that is superimposed on a background of other challenges.

### Cardiovascular disease

Cardiovascular disease (CVD), the leading COD worldwide (Heron 2015), has properties that cast it into the class of extrinsic mortality. Firstly, CVD exhibits an exponentially increasing mortality rate with age^5^. The distal and proximal factors have clear extrinsic mortality character. For example, atherosclerosis, the accumulation of plaque within the artery wall, becomes the distal factor of CVD death while the immediate cause of death, e.g. myocardial infractions, become the proximal factor. Looking further, atherosclerosis is an inflammatory disease promoted by inflammaging and immunosenescence (Epstein and Ross 1999; Legein et al. 2013). A surrogate measure of atherosclerosis, the carotid intima-media thickness, has a log-linear relationship with cardiovascular events (Bots et al. 2016) and studies suggest the thickness increases approximately linearly with age (Polak et al. 2011). Furthermore, telomere length, which is associated with the progression of atherosclerosis, also has a linear relationship with age (Aviv et al. 2015; Haycock et al. 2014). The linear properties are all captured by the decline of vitality with age. For the proximal factor, myocardial infractions are triggered by factors such as physical activity, emotional stress, sexual activity and eating (Čulić et al. 2005), which together can be represented by probability distributions of challenge magnitude and frequency.

Importantly, the distributions of challenges for CVD and ID are significantly different. CVD challenges have high frequency, essentially occurring daily, but their magnitudes are low. In contrast, ID challenges occur with low frequency, on the order of years, but their magnitudes are high. These differences should result in ID, when of sufficient frequency, masking the contribution of CVD to extrinsic mortality. This masking effect is discussed in the section on challenge masking.

### Cancer

Malignant neoplasms currently represent the second major COD worldwide (Heron 2015). For many types of cancer the rate of mortality increases approximately exponentially through early-to-middle old age (60 to 80 years) and then declines (Bonafè et al. 2001; Ukraintseva and Yashin 2003)^6^. This nonlinear pattern might be explained by cancer’s multistage etiology in which DNA mutations to stem cells promote their clonal growth that then can transition into senescence or a malignant neoplasm (Hanahan and Weinberg 2011). The progression of cancer, like CVD, involves immunosenescence and in particular accumulation of senescent cells with age is thought to promote tumor growth (Campisi 2013; Vasto et al. 2009). Therefore, the stochastic property of vitality provides a framework that may capture the age evolving properties of cancer development. However, the decline in the cancer mortality rate in old age requires further discussion.

Two explanations for the decline of cancer in old age have been proposed. One involves selective removal of frail individuals that are genetically predisposed to cancer, leaving cancer-resistant individuals in old age (Vaupel and Yashin 1986). This mechanism assumes genetic heterogeneity in the inflammation response in which individuals with a weaker response to inflammations are presumably more resistant to cancer (Vasto et al. 2009). This process might in part be captured in the vitality framework by the mortality rate plateau produced by the absorption of stochastic vitality paths into the zero boundary (Weitz and Fraser 2001). However, this mechanism does not explain strong declines in cancer incidence in old age. A reduced inflammation response could also result from remodeling of the immune system without invoking population heterogeneity (Bonafè et al. 2001; Ukraintseva and Yashin 2003). Additionally, decline in cancer incidence may also involve telomeres through a protective effect of telomere attrition on senescence (Blackburn et al. 2015; Collado et al. 2007), which could inhibit clonal growth. These mechanisms are not mutually exclusive.

The salient point here is that the distal factor of cancer involves greater complexity than what is captured by the current two-process vitality model. In any case, cancer does not fit well into a challenge paradigm of extrinsic mortality because the mutations that initiate clonal growth and cancer can result from intrinsic or extrinsic challenges (Tomasetti and Vogelstein 2015; Wu et al. 2016). Additionally, the latency between an initial mutation and mortality can span years, which precludes a clear characterization of a specific challenge frequency or magnitude for the process. The proximal causes of cancer deaths are more characteristic of the boundary absorption process underlying intrinsic mortality. For example, exhaustion of survival capacity is readily represented by cachexia, a metabolic process with extreme weight loss that is the immediate cause of 20% of cancer deaths (Argiles et al. 2014). Because of the boundary absorption nature of the proximal factor of cancer and the indeterminacy of the source of cancer initiation we surmise that the vitality model imperfectly captures cancers as intrinsic mortality. However, it is likely that intrinsic mortality of the model also is driven in part by CVD-like diseases through challenge masking discussed below.

## Model

### Equations

As discussed in the introduction the two-process model has its foundations in the vitality challenge framework of Strehler and Mildvan (1960) and borrows extrinsic-intrinsic mortality terminology of Carnes et al. (1996). However, in focusing on processes the model of Li and Anderson (2013) provides a more generalized classification in which mortality involves distinct distal and proximal factors. For the extrinsic mortality component, the distal factor is represented by age-declining deterministic vitality and the proximal factor is represented by random challenges. For intrinsic mortality, the distal factor is represented by age-declining stochastic vitality and the proximal factor is represented by boundary absorption (Li and Anderson 2013), which can also be viewed as weak magnitude challenges. To this base model we now include juvenile mortality with a distal factor represented by age-increasing juvenile vitality and a proximal factor represented by juvenile challenges. Assuming challenges to the respective vitalities do not affect the distributions of the other vitalities then the mortality rates are independent and the total mortality rate is

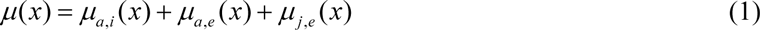

where subscript (*a, i*) denotes the adult intrinsic process and (*a, e*) and (*j, e*) denote the adult and juvenile extrinsic processes respectively. The interactions of vitalities and challenges are depicted in Fig. 1 and described below.

**Fig. 1.**
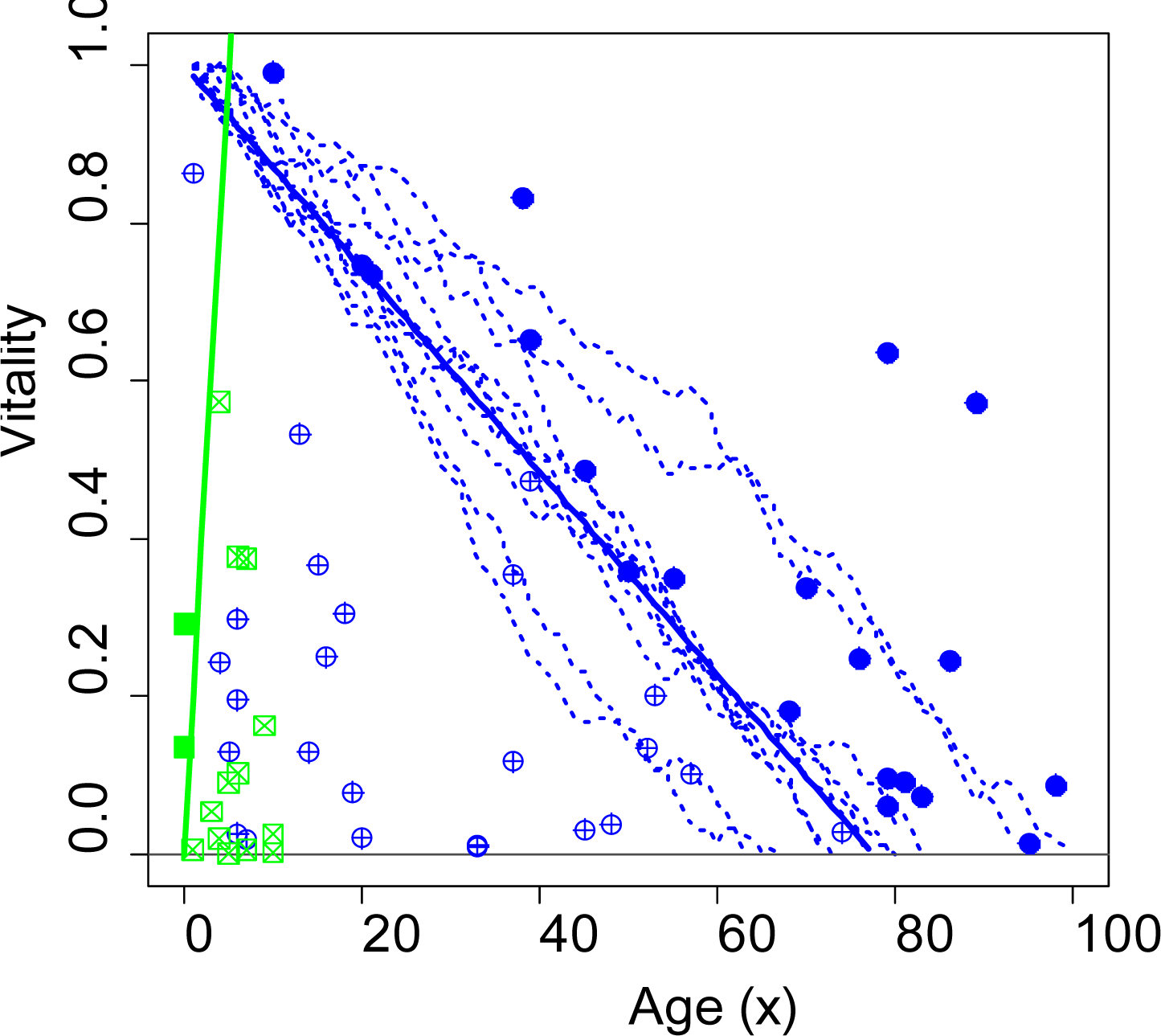
Depiction of extrinsic and intrinsic mortality processes for equation (5). The green and blue lines represent population-level measures of juvenile and adult vitality respectively where from birth juvenile vitality increases from zero and adult vitality decreases from one. Green squares and blue circles denote juvenile and adult vitality extrinsic challenges. Filled symbols denote juvenile and adult extrinsic deaths in which the challenges exceed juvenile and adult population-level vitalities. Correspondingly, unfilled symbols represent challenges not exceeding the reference vitalities. Adult intrinsic mortality is represented by individual stochastic vitality paths intersecting the zero boundary which we denote an intrinsic challenge. The figure was generated using *r* = 0.0129, *s* = 0.0126, *λ* = 0.045, *β* = 0.40, *γ* = 0.15 and *α* = 1 corresponding to the fit to period data for Swedish females in 1900

#### Adult intrinsic mortality

Adult intrinsic mortality is generated by boundary passage of a Wiener process (Anderson 2000; Li and Anderson 2013; Weitz and Fraser 2001) as 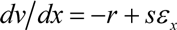 where *r* is the mean rate of loss of vitality, *s* is the stochastic intensity of the loss rate and *ε*_*x*_ is a white noise process. The probability distribution of the first passage time of stochastic vitality trajectories from a unit initial vitality at age *x* = 0 to the death boundary *v* = 0 (Fig. 1) is given by the inverse Gaussian distribution (Chhikara and Folks 1989), 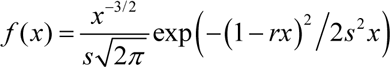. Survival from adult intrinsic mortality only, is 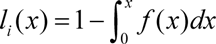 and the adult intrinsic mortality rate is

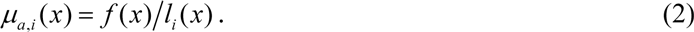

#### Adult extrinsic mortality

Adult extrinsic mortality, following (Li and Anderson 2013; Strehler and Mildvan 1960), is generated by environmental challenges to adult vitality. Challenge frequency has a Poisson distribution with mean *λ*. The challenge magnitude *z* has an exponential distribution 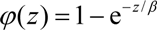 with a mean *β*. The conditional extrinsic mortality rate is *m*_*e*_ 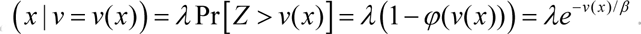. Integrating over vitality states, the population-level adult extrinsic mortality rate at age *x* is 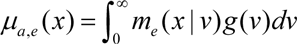 where the age-dependent distribution of vitality *g*(*v*) depends on both the loss of stochastic vitality through absorption into the zero boundary and its modification by the preferential elimination of low-vitality individuals via challenges. Because *g*(*v*) has no closed form, age-dependent vitality is approximated as a deterministic function as 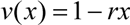 giving the adult extrinsic mortality rate (Li and Anderson 2013)

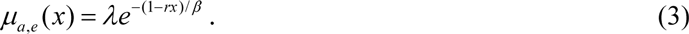

Note in (Eq. (3) mortality results from challenges to a linear approximation of the age-declining stochastic vitality distribution. The extrinsic mortality rate is equivalent to the Strehler and Mildvan (1960) interpretation of the Gompertz (1825) mortality model 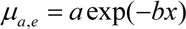 such that the Gompertz coefficients are related to the extrinsic mortality coefficients as 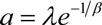 and 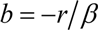.

#### Juvenile extrinsic mortality

Juvenile extrinsic mortality is generated by juvenile challenges to deterministically increasing juvenile vitality. The development follows that of Eq. (3) except at age *x* = 0 juvenile vitality is zero and it increases giving *v*′(*x*)=*r*′*x*, where the prime designates juvenile parameters equivalent to the adult parameters. The juvenile extrinsic mortality rate from juvenile challenges is then

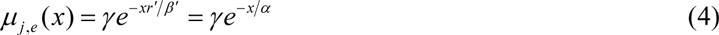

where *γ* is a base juvenile mortality rate and *α* = *β*′/*r*′characterizes the duration of the juvenile stage.

#### Total mortality

The total mortality rate combines Eqs. (2–4) into Eq. (1) giving

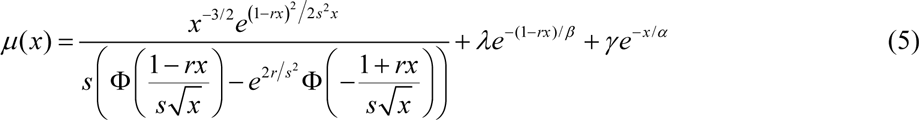

where Φ is the cumulative standard normal distribution 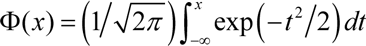. Equation (5) parameters are estimated by fitting survivorship curves or mortality rate data with a maximum-likelihood fitting routine (Salinger et al. 2003). A family of vitality models in R code is available at CRAN.R-project.org/package=vitality. Equation (5) is fit with the package function vitality.6p.

### Model properties

The model partitions three types of mortality, which are defined by their distal and proximal factors. However, there is no absolute definition of the partitions, nor one-to-one relationships with COD. Thus, to address these indeterminacies sections below briefly discuss the properties of intrinsic and extrinsic mortalities and their hypothesized relationships to COD and physiological processes at the cell and system levels. The third section discusses the issue of resolving proximal factors of mortality.

#### Intrinsic mortality

For intrinsic mortality, we assume the distal factor represents the stochastic loss of immune system capacity through immunosenescence and the proximal factor, mathematically the passage of vitality into the zero boundary, represents death by the inability of the immune system to maintain homeostasis. Cancer is largely captured as intrinsic mortality because the endpoint is typically a wasting process, e.g. cachexia. Additionally, the patterns of log mortality rate versus age are similar for cancer and intrinsic mortality, i.e. for both, the increase of mortality rate slows down with age. The model generates this pattern by the removal of lower vitality (cancer susceptible) individuals at earlier ages leaving the higher vitality (cancer resistant) individuals in older ages. Mortality associated with CVD may also be represented as intrinsic mortality under challenge masking, which is discussed below. However, the current model does not capture the reduction of cancer rate in very old age.

#### Extrinsic mortality

For adult extrinsic mortality we assume the distal factor represents the mean rate of decline of immune system capacity through immunosenescence and the proximal factor represents extrinsic challenges to the system’s capacity. For juvenile extrinsic mortality we assume distal vitality represents a linear increase in the immune system strength in early life and the proximal factor represents challenges to the immune system. The increase of juvenile vitality has no effect on the decrease in adult vitality, but for adults, extrinsic and intrinsic mortalities are linked since both are assumed to depend on the rate of immunosenescence.

#### Challenge masking

The distributions of adult challenge magnitude (*β*) and frequency (*λ*) do not fully capture the range of patterns expected by ID and CVD. ID challenges tend to be strong and infrequent, e.g. typhoid epidemic, while CVD triggers are weak and frequent, e.g. physical exertion. While both types co-occur, the model estimates properties of high magnitude challenges when their frequency is above some threshold level and when it drops below the threshold the model estimates the properties of the low magnitude challenges (Li and Anderson 2013). We denote this property “challenge masking” in that strong challenges mask the effect of weaker ones resulting in the weaker challenges being represented as vitality boundary absorption when they co-occur with strong challenges. In essence, weak challenges (e.g. CVD triggers) in the presence of strong challenges (e.g. ID events) in part can be shifted to the intrinsic mortality category but in the absence of the stronger challenges they contribute mainly to the extrinsic mortality category. In addition, unmasking of weaker challenges also involves an actual decline in the extrinsic mortality rate because ID challenges are removed. Ultimately, the clarity of the intrinsic and extrinsic mortality partition depends on properties of the challenges. An important point is that COD categories included in the intrinsic mortality can depend on the extrinsic challenges, which change over an epidemiological transition.

## Results

### Mortality rate patterns with age

Figure 2, showing patterns of the three mortality processes for Swedish period mortality data^7^, illustrates the model interpretation of changes in processes underlying the doubling of life expectancy (~40 to ~80 years) between the 19^th^ and 20^th^ centuries (Sundin and Willner 2007). This lifespan doubling is explained by reductions in juvenile mortality and adult extrinsic mortality for ages < 70 years. However, for ages > 70 years extrinsic mortality rate increases while the intrinsic mortality rate decreases. We attribute this cross-century change to two processes. First, the ID (high magnitude/low frequency) challenges diminish, which reduces juvenile and adult extrinsic mortalities. Second, the reduction of ID challenge frequency unmasks the CVD challenges such that the model shifts CVD-type mortalities from intrinsic to extrinsic categories. Thus, the modeled intrinsic mortality rate decline across the centuries reflects a decline in ID-type mortality plus an apparent decline resulting from a shift of CVD-type mortalities from intrinsic to extrinsic categories. The dominance of extrinsic mortality in old age after the epidemiological transition comports with CVD now being the leading COD. Correspondingly cancer, which the model associates with intrinsic mortality, is the second leading COD (Heron 2015). The decline in childhood mortality from the 19^th^ to the 20^th^ century is postulated to be from both a reduction in childhood disease challenges and an increase in the rate of immune system development, plausibly related to increased nutrition in the 20^th^ century (Köhler 1991). The current model does not include young adult challenges, which is evident by the rise of the mortality rate above the extrinsic mortality line between ages 10 and 25 years (Fig. 2).

**Fig. 2.**
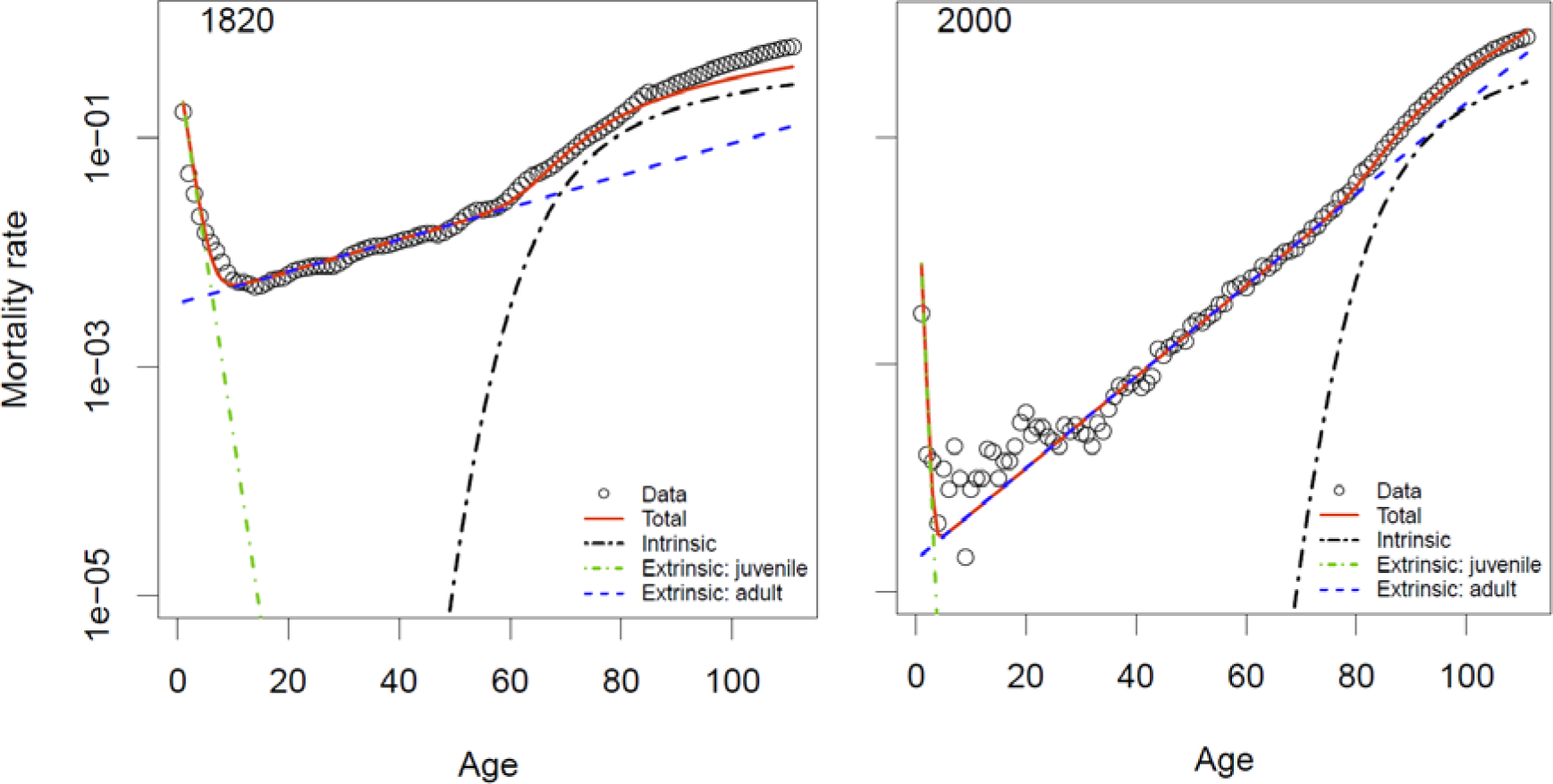
Period mortality rate components vs. age for Swedish female period data for 1820 and 2000. Data depicted by circles. Modeled total mortality rate is depicted by red lines. Green, dark blue and light blue lines depict mortality rate components adult intrinsic (*µ*_*i*_), adult extrinsic (*µ*_*e,a*_), and juvenile extrinsic (*µ*_*e,j*_) respectively. Fitted parameters are *r* = 0.0129, *s* = 0.0126, *λ* = 0.045, *β* = 0.400, *γ* = 0.2033, *α* = 0.75 for year 1820 and *r* = 0.0107, *s* = 0.0077, *λ* = 0.123, *β* = 0.115, *γ* = 0.0076, *α* = 2.56 for year 2000

#### Juvenile extrinsic mortality

The juvenile extrinsic mortality rate, characterized by straight lines in the log mortality rate graph (Fig. 3*a*), clearly shows two clusters. One cluster (red-green lines, 1751-1905) has a shallow slope with age and the other cluster (purple lines, 1960-2010) has a steeper slope. The epidemiological transition (1905-1960) is evident as a fan of blue lines between the two clusters. The change in the juvenile extrinsic mortality rate exhibits a progressive decline in both the initial rate *γ*, characterized by line intercepts at age 0, and the duration of juvenile mortality *α*, characterized by line slopes. The parameter *γ* reflects both neonatal mortality and the frequency of challenges experienced by juveniles. The crossover age, at which the juvenile and adult extrinsic rates are equal, is depicted by circles passing through each respective line. In the early period (red-green cluster) the crossover age varies from ~ 5 to 7 years of age, mostly as a result of changes in line slopes, which are expressed by *α*. We suggest this variability and trend in the crossover age was driven by a series of famines superimposed on a gradual improvement in the nutrition status of children. This is elaborated below in discussion of Fig. 5*e*. The further reduction of the crossover age and decline in crossover age mortality rate is driven by the reduction in the adult extrinsic mortality rate as characterized by *r* (Fig. 5*a*).

**Fig. 3.**
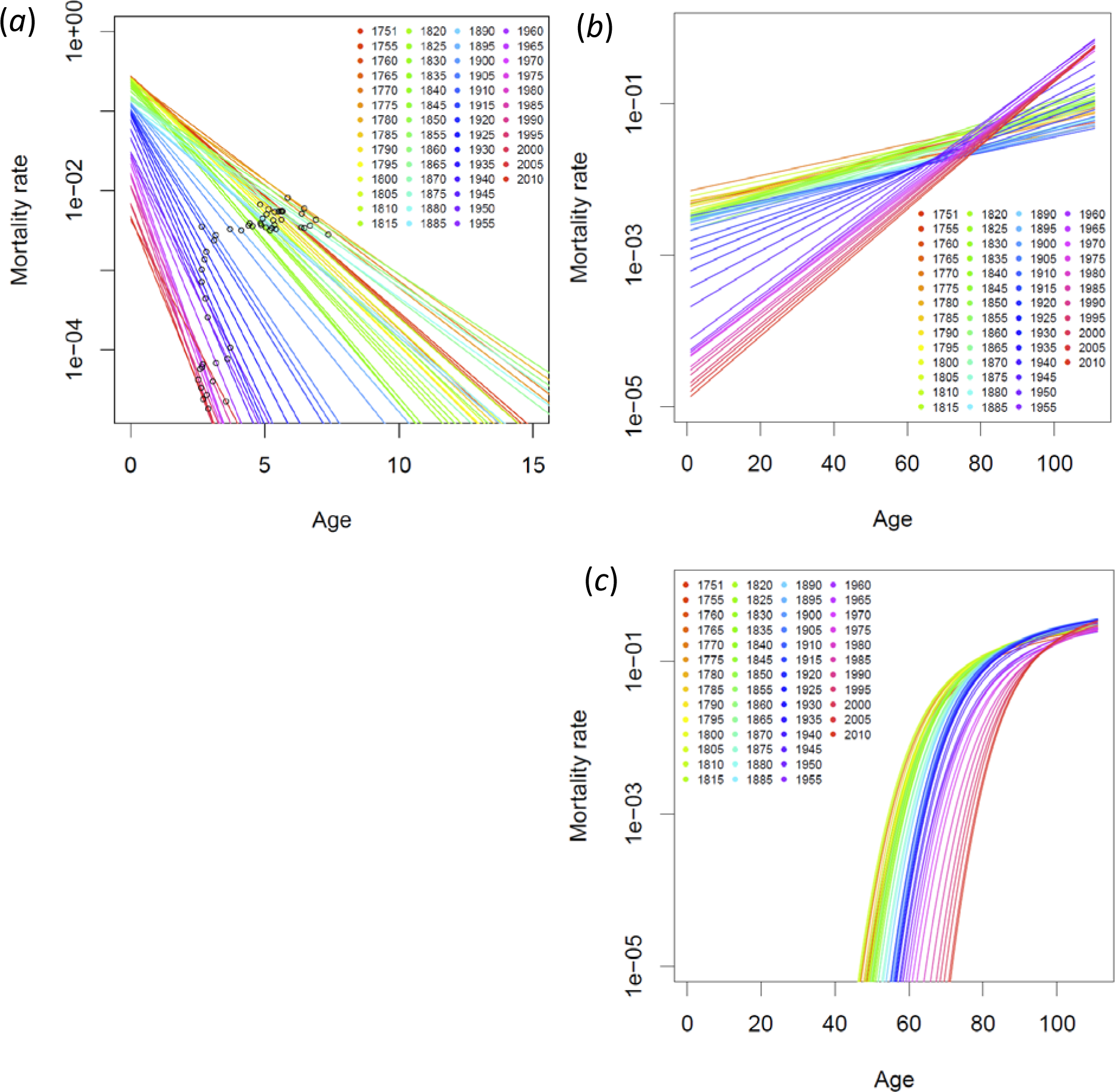
Lines depict mortality rate components (year^−1^) in equation 5 fit to Swedish female period data grouped in 5-year increments (1751-2010). In general, for each graph the mortality rate components decline for increasing 5-year intervals: (*a*) Juvenile extrinsic mortality rate. Circles indicate cross over ages where juvenile and adult extrinsic mortality rates are equal. (*b*) Adult extrinsic mortality rate. (*c*) Adult intrinsic mortality rate.

#### Adult extrinsic mortality

The pattern of adult extrinsic mortality rate lines (Fig. 3*b*) cluster in the same manner as the juvenile lines (Fig. 3*a*). Within each cluster the lines progressively, and uniformly, shift downward over years as a result of the gradual reduction across all ages in the extrinsic mortality rate. The two clusters intersect at approximately age 75 years, indicating that in this interval the estimated extrinsic mortality was approximately constant across time. Thus, for ages below 75 years, the extrinsic mortality rate declined over time due to a reduction of strong ID challenges, while above 75 years the rate increased because of the corresponding unmasking of the low magnitude CVD-type challenges. In the red-green cluster, extrinsic mortality was driven mainly by ID challenges, which occurred randomly over the lifespan, thus the rate of mortality from ID challenges was similar for young and old individuals resulting in a shallow slope in the cluster. In contrast, the purple cluster extrinsic mortality was dominated by low magnitude challenges, which occurred later in life, resulting in a steeper slope for the cluster and a strong age dependence in extrinsic mortality.

#### Adult intrinsic mortality

The pattern of intrinsic mortality completes the picture of longitudinal changes (Fig. 3*c*). Notably, intrinsic mortality in old age (> 95 years) changed little over the years suggesting the rate of mortality from cancer in very old age has changed little over time. However, the across-decade changes in intrinsic mortality are quite significant for middle and early old age. For example, at age 80 the intrinsic mortality rate was ~0.1 in year 1750 and 0.001 in year 2010; a two order of magnitude decline in intrinsic mortality. This decline is not the result of the challenge masking effect since essentially the magnitude of decline occurred within the red cluster (years 1960-2010) in which the extrinsic challenge properties had stabilized. The cause of this decline reflects the cumulative changes over the lifespan of the individuals, as expressed by *r* (Fig. 5a), and likely reflects improved nutrition as well as the reduction of exposure to illness through better health programs.

#### Uncertainty in rates

Figures 2 and 3 demonstrate significant and continuous differences in the intrinsic and extrinsic mortality rates across two centuries. Figure 4*a, b* illustrating the density distribution of mortality rates estimates reveals that temporal changes in rates are much greater than the uncertainty of estimates based on survival patterns over five-year intervals. Extrinsic mortality exhibits a clear progression of declining rates at age 40 with a narrower distribution of mortality rates with each time interval. The pattern of the intrinsic rate is similar with a general trend toward lower mortality rates at age 85. While extrinsic rate progressed consistently year after year, most movement in the intrinsic rate began after ~1940, which is consistent with our findings in (Sharrow and Anderson 2016a).

**Fig. 4.**
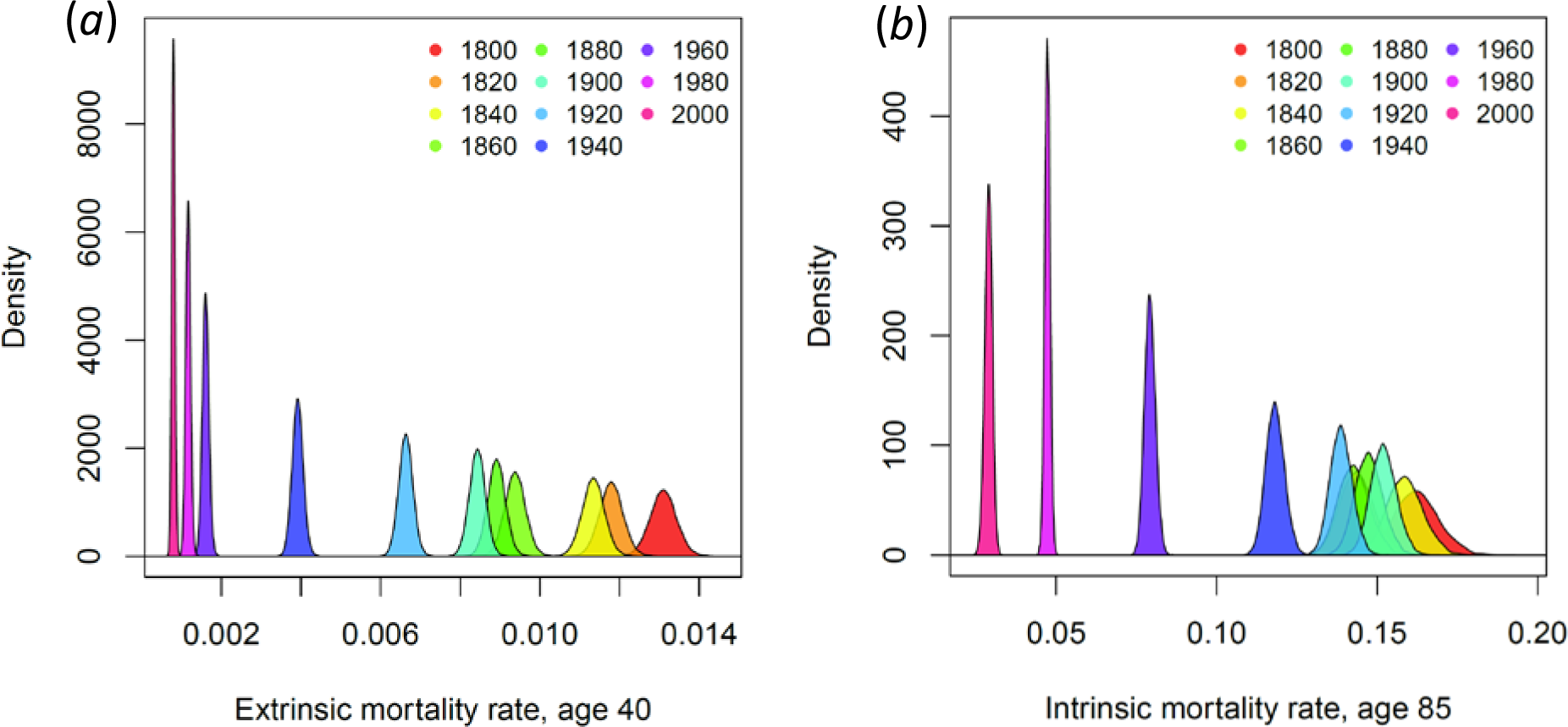
Depiction of the distribution of a) extrinsic mortality rates at age 40 and b) intrinsic mortality rate at age 85 for every 20 years (based on 5-year life tables, e.g. 1800-1805,). The distributions are generated by calculating mortality rates with a distribution of vitality parameters generated by sampling from a normal distribution with mean equal to the maximum likelihood parameter estimate and standard deviation equal to the parameter standard error generated from the vitality fitting package with the initial population set to 100,000.

### Vitality parameter longitudinal patterns

Figure 5 depicts the patterns of the vitality model parameters for male and female Swedish populations over two centuries. Besides a general improvement in conditions promoting a doubling of longevity over this period of time, the parameters reveal the complexity in the changes that occurred over decades as well as the complexity of the epidemiological transition. Because of the effect of challenge masking involved with the epidemiological transition we discuss the periods before and after the transition separately.

#### Before mid-20^th^ century

Before the mid-20^th^ century, adult extrinsic mortality declined slowly and erratically (Fig. 3*b*) as a result of a slow steady decline in the rate of loss of vitality (Fig. 5*a*) and variability in the challenge magnitude and frequency (Fig. 5*c* and 5*d*). The gradual decline in *r* in the 19^th^ century (Fig. 5*a*) corresponds with a significant increase in agricultural production beginning 1790 (Olsson and Svensson 2010), which was driven by the replacement of small farms with large capitalistic farms (Möller 1990). The link between grain production and *r* is supported by an observed correlation between the price of rye and the mortality rate over the first half of the 19^th^ century. A 20% increase in grain price corresponded with a 5-6% increase in mortality the next year (Dribe et al. 2012). Because both food availability and its price would affect the level of nutrition (Schaible and Kaufmann 2007), we postulate that the pattern of *r* over time reflects the effects of nutrition on the rate of development of immunosenescence, i.e. higher values of *r* indicate faster rates of immunosenescence.

**Fig. 5.**
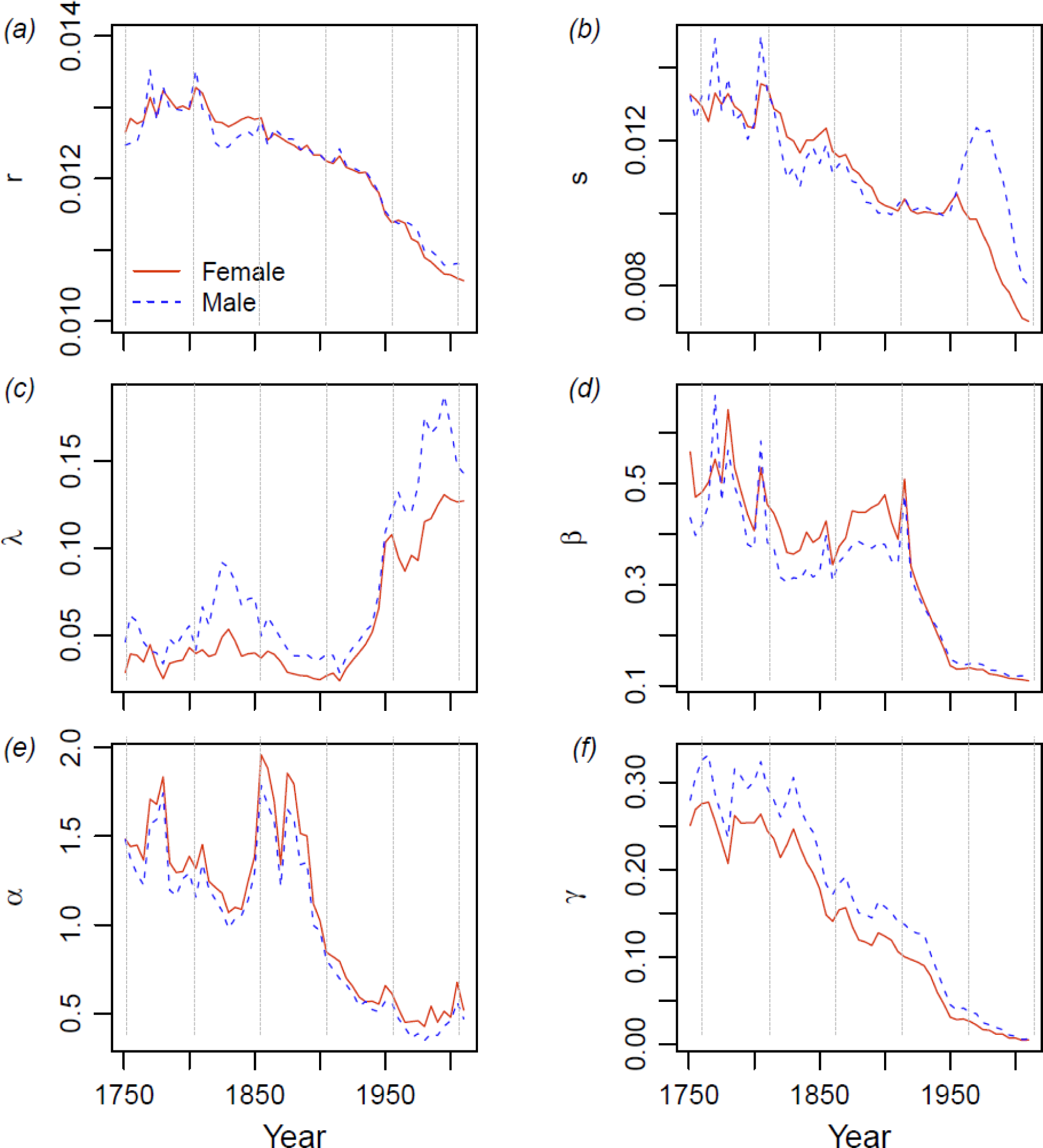
Longitudinal patterns of vitality parameters for the Swedish population (1750-2010). Male and female patterns are depicted as dashed (blue) and solid (red) lines separately. (*a*) vitality loss rate (year^−1^), (*b*) variation of vitality loss rate (year^−1/2^), (*c*) average adult extrinsic challenge frequency (year^−1^), (*d*) average adult extrinsic challenge intensity, (*e*) half-life of juvenile mortality (year), (*f*) base juvenile mortality rate (year^−1^)

Vitality rate variability, *s*, was high in the 18^th^ century and then declined over the 19^th^ century (Fig. 5*b*). This term characterizes the degree of rectangularization in the survival curve, i.e. the steepening of slope of the survival curve about the mean age of life expectancy (Li and Anderson 2009). It also quantifies the heterogeneity in the intrinsic mortality rate of the population. The high levels of *s* in the 18^th^ and early19^th^ century likely correspond with the differential in mortality between the rural and urban populations (Edvinsson and Lindkvist 2011; Schumann et al. 2013). Nutrition levels were higher and infection levels were lower in the rural environment. Also early in the 19^th^ century mortality rates were lower in the higher social economic groups than in the lower groups (Bengtsson and Dribe 2011). The differential in mortality by both measures declined steadily through the 19^th^ century and into the first part of the 20^th^ century (Bengtsson and Dribe 2011). Thus, we suggest *s*, is potentially a quantitative measure of the effects of heterogeneity in population nutrition on mortality during this period.

Noteworthy, in Fig. 5*c* the frequency of infectious disease and physical injury challenges, *λ*, did not significantly change through the 18^th^ and 19^th^ centuries for females. However, the challenge frequency increased significantly for males in the first half of the 19^th^ century. This peak in *λ* corresponds with excessive male mortality between 1820-1850, which was postulated to be caused by increased alcohol consumption in males (Sundin and Willner 2007). Also, the male versus female differences in challenge frequency (Fig. 5*c*) comport with gender-linked risk taking behavior, which was about 20% lower in females than in males (Byrnes et al. 1999). Challenge magnitude (Fig. 5*d*) exhibited distinct spikes for the years 1808 and 1918, which correspond with disease during the Finnish war (Mielke and Pitkänen 1989) and the influenza pandemic (Sundin and Willner 2007) respectively, suggesting that *β* quantifies the virulence of the environment. This signal in *β* corresponds with the effects of pandemics to temporarily shift mortality to younger age groups (Reichert et al. 2012). Finally, the higher magnitude *β* in females comports with the allocation of resources to males over females (Klasen 1998) in the 18^th^ and 19^th^ centuries. Thus, the extrinsic challenge parameters *λ* and *β* quantify notable environmental-and behavioral-associated changes in health.

The index of juvenile mortality duration, *α*, exhibited peaks (Fig. 5*e*) corresponding with famines in 1772-1773 and ~ 1860-1870 (Sundin and Willner 2007). Note that starvation itself represents only a small fraction of the mortality associated with famine. The majority of deaths are due to infectious diseases associated with malnutrition (Sundin and Willner 2007). Thus, we suggest that because *α* = *β*′/r′ it follows that the peaks in *α* were the result of reduced food supply that decreased the rate of immune system development, i.e. reduced *r*′. A series of nine cholera epidemics between 1834 and 1874 likely contributed to the elevated *α* over the same period through the effect of cholera on the juvenile challenge magnitude *β*′. The steady decline in the initial mortality rate *γ*(Fig. 5*f*) through the mid-20^th^ century does not correspond with specific health events. Noteworthy, the initial mortality rate was uniformly higher in boys than girls, a feature also observed in other populations (Sundin and Willner 2007).

#### After mid-20^th^ century

The mid-20^th^ century epidemiological transition is evident by a sharp decline in the vitality loss rate, *r*, in males and females about ~1945 (Fig. 5*a*). About 1950 the trend in female *s* exhibited a decline while the male trend increased, peaking ~1975 (Fig. 5*b*). The decline in female *s* is potentially a continuation of the postulated trend in decreasing heterogeneity in the rate of immunosenescence in the female population. The rise and peak in *s* for males can be explained by increasing heterogeneity in their rate of immunosenescence due to a divergence in the social economic pattern of smoking. About 1950 the male smoking rate started to decline in the professional/managerial group but the decline in the agricultural/industrial group did not begin until the end of the century (Diderichsen and Hallqvist 1997). Thus, the peak in *s* in ~1975 likely quantifies the lag in the reduction of smoking between the two groups. The model does not identify the COD associated with the peak in *s*. However, in the period 1965-80 the leading COD in the agricultural/industrial group was CVD (Diderichsen and Hallqvist 1997). Because *s* only occurs in the intrinsic mortality (Eq. 2) and its level is most influenced by the curvature of the mortality rate in old age, the peak was driven by differences in the mortality rates of the two social economic groups in old age.

The rapid decline in challenge magnitude from 1920 to 1950 (Fig. 5*d*) is coincident with measures to control tuberculosis, the discovery of streptomycin (Daniel 2006) and development of a range of antibiotics for other infectious diseases. The slower decline in challenge magnitude post-1950 can be explained by two factors. Firstly, the extrinsic mortality in this period was driven by CVD-type mortalities, which were flat over the period (Vangen-Lønne et al. 2015). Secondly, CVD mortality is triggered by every-day physical exercise which has not changed significantly over the last half century. Thus, the flat pattern of *β* reflects the low magnitude of proximal triggers associated with CVD mortalities. Finally, in the 21^st^ century extrinsic mortality exceeds intrinsic mortality, which comports with the COD data in which in the 21^st^ century mortality from degenerative diseases exceeds that from malignant neoplasms (World Health Organization 2010).

The 20^th^ century decline in *α* (Fig. 5*e*) corresponds with improvements in Swedish postnatal care and child nutrition (Köhler 1991). The decline in the initial juvenile mortality rate *γ* likely reflects improvements in neonatal care in the second half of the century (Fig. 5*f*).

## Discussion

This paper describes a process POV approach to modeling mortality patterns and outlines a framework for characterizing COD in terms of separate processes leading up to death and its immediate cause. We designate these the distal and proximal factors of mortality. The framework has its basis in the Strehler-Mildvan (1960) interpretation of the Gompertz (1825) model that characterizes interactions in terms of external challenges (proximal factor) exceeding the level of the age-declining vitality (distal factor). This framework was extended in Li and Anderson (2013) by including a second mortality process, characterized by the absorption of vitality into a zero boundary. This two-process framework defines two distinct forms of mortality: extrinsic mortality induced by challenges to deterministic vitality and intrinsic mortality induced by absorption of stochastic vitality. Notably, the partition has different properties to the extrinsic-intrinsic partition proposed by Carnes and colleagues (Carnes and Olshansky 1997; Carnes et al. 1996).

We extend the process POV by seeking candidate biological processes underlying the distal and proximal factors. We first reinterpret vitality absorption as functionally equivalent to low magnitude-high frequency challenges exceeding low level vitality. In this revision, intrinsic and extrinsic mortalities then both have distal factors that measure vitality and proximal factors that define challenges. Using this distal/proximal framework, we next define juvenile mortality as challenges exceeding age-increasing juvenile vitality. The resulting three mortality rates are define by model parameters characterizing the dynamics of the distal and proximal factors: intrinsic mortality depends on the mean and variance in the rate of loss of vitality (*r, s*), extrinsic mortality depends on *r* and the mean magnitude and frequency of challenges (*β*, *λ*) and juvenile extrinsic mortality depends on the base juvenile mortality rate and duration of the juvenile stage (*γ*, *α*) where *α* = *β*′/*r*′.

The model fit well the Swedish survivorship curves with juvenile extrinsic mortality dominating early ages, adult extrinsic mortality dominating middle ages and intrinsic mortality dominating old ages. The resulting mortality rates and distal/proximal factors also exhibit clear patterns that comport with environmental and social-economic changes between the 19^th^ and 21^st^ centuries.

We explored possible biological processes underlying the patterns by first reviewing recent literature on cellular hallmarks of aging and age-dependent changes in physiological systems. Although a multitude of processes contribute to aging, two findings standout relative to understanding vitality. Firstly, evidence suggests biochemical pathways to mortality merge in a central pathway that can be described by a single variable (Stroustrup et al. 2016). Secondly, the immune system, with its functions of eliminating pathogens and repairing tissue is involved with virtually all aging processes. From these findings and others, we considered the 20^th^ century epidemiological transition under the assumption that vitality parameters reflect the historical pattern of population immune system health. The exercise revealed plausible and thought-provoking connections between environment, health and mortality.

However, the exercise also illustrated the limitations of the mortality partitions in quantifying categories of COD. While the model parameters and historical information exhibited clear and biologically meaningful correspondences, the partition of mortality did not have a one-to-one correspondence with COD. Specifically, prior to the 20^th^ century epidemiological transition, adult extrinsic mortality tracked patterns of infectious disease and adult intrinsic mortality largely tracked degenerative diseases and malignant neoplasms. With the reduction of infectious disease during the epidemiological transition, the extrinsic mortality largely tracked the pattern of degenerative diseases while the intrinsic mortality appeared to capture mostly malignant neoplasms. We conclude that the partition of mortality components depends on the characteristics of the challenges. We termed this “challenge masking” to reflect the effect of strong challenges (e.g. typhoid) masking the effects of weak challenges (e.g. stress induced myocardial infarctions). This masking was not unexpected and illustrates that while the model quantitatively partitions mortalities, the processes underlying the partition, and in particular specific causes of death, are not easily separated with mortality data alone.

In a practical sense, we view mortality partitions, such as the adult extrinsic/intrinsic partition, as hypotheses of the contributions of different competing distal and proximal factors shaping COD patterns. Validating these hypotheses and further identifying the underlying mechanisms will require independent quantitative and qualitative information. For example, the model prediction of extrinsic mortality dominating intrinsic mortality in old age (Fig. 2*b*) is supported by data showing that CVD is the leading cause of death while cancers are the second leading cause. This ordering of causes supports the model inference that CVD is a form of extrinsic mortality while cancers are a form of intrinsic mortality. Furthermore, the model provides a quantitative assessment of the change in significance of CVD and cancer with age and in particular characterizes the reduction of cancer incidence relative to CVD in very old age.

It remains to be determined to what degree mortality partitions can be better linked to specific causes of disease and underlying biological processes. Some extensions should be relatively straightforward. For example, characterizing excess mortality in young adults with a distribution of challenges representing risk-taking fits with theories and observations of the young adult mortality hump. However, partitioning contributions of CVD and cancers to patterns of old age mortality is more problematic. In the current model form these processes share a common distal factor, the rate of loss of vitality *r*. However, the etiologies of the two diseases are significantly different and therefore a more precise partition will likely require separate rates of loss of vitality for CVD and forms of cancer. In any case, irrespective of the complexities, the process POV offers a new approach to reconcile patterns of cause-specific mortalities with patterns of all-cause mortality, a topic under active discussion (Alai et al. 2015; Beard 1971; Foreman et al. 2012; Tuljapurkar 1998).

Lastly, we return to the value of the process POV, which in spite of its long history, currently makes only a small contribution to population sciences. Our perspective of the value of the process approach can be illustrated through the debate on the limits of human longevity. Over the years, several papers have predicted a plateau in longevity based on largely qualitative assumptions that gains over the 20^th^ century are unlikely to be repeated with further life-style changes and medical enhancements (Olshansky et al. 2001). A counter argument, that a plateau is not impending, is based on the fact that life expectancy has increased in a linear manner for the past 160 years. Therefore, unforeseen future improvements may very much increase longevity in the 21^st^ century (Oeppen and Vaupel 2002). Counter to this, intrinsic and extrinsic partitions of lifespan may be reaching an asymptote (Sharrow and Anderson 2016a) and maximum lifespan is not increasing (Dong et al. 2016). However, none of these arguments are based on an understanding of what processes contributed to the past trends in longevity nor what changes are possible at the cellular level that ultimately determines longevity. It seems clear that merging the sciences that study cellular mechanisms of aging and mortality with the sciences that project consequences to populations is the next logical step in understanding the limits of human lifespan. We suggest this step requires a process POV framework.

1 We do note that another significant body of theory links biological processes to mortality but this framework links biological processes directly to the mortality rate and does not track the cumulative effects of processes leading up to mortality (e.g. Yashin et al. 2012; Yashin et al. 2016; Yashin et al. 2000).

2 The idea of partitioning mortality into intrinsic and extrinsic parts was originally proposed by Carnes and colleagues (Carnes and Olshansky 1997; Carnes et al. 1996). They defined extrinsic mortality as avoidable mortality and intrinsic mortality as unavoidable mortality. The partition was demonstrated using COD data to calculate extrinsic mortality and intrinsic mortality was calculated as the difference between extrinsic and all-cause mortality (Carnes et al. 2006). However, the partitions based on the two vitality processes (Li and Anderson 2013) and on COD yield different mortality patterns with age. For both partitions, extrinsic mortality dominates in younger ages but the intrinsic mortality patterns are significantly different in the two methods. In the COD method intrinsic mortality has a Gompertz-like age distribution while in the vitality method intrinsic mortality is negligible in young age, then rises rapidly and plateaus in old age (Weitz and Fraser 2001). Consequently, the two methods of partitioning mortality cannot be easily compared.

3 Other terms for vitality include viability (Weitz and Fraser 2001), vital capacity (Whitmore 1986) and whole organisms state variable (Stroustrup et al. 2016). Other frameworks have proposed similar ideas, e.g. frailty, a fixed quantity endowed at the beginning of life (Vaupel et al. 1979) and redundancy theory (Gavrilov and Gavrilova 2003), but these applications were based more in a mortality rate POV and are not explored here.

4 Carnes et al. (2008) based on (Arking 2006) suggested five characteristics of aging denoted C.U.P.I.D as Cumulative, Universal, Progressive, Intrinsic and Detrimental.

5 Example age distribution of the mortality from cardiovascular diseases for the female Swedish population 2000-2004 is available from the Institute for Health Metrics and Evaluation at ihmeuw.org/3tzc.

6 Example age distribution of incidence of colon and rectum cancer for male Swedish population 2000-2009 is available from the Institute for Health Metrics and Evaluation at ihmeuw.org/3v13.

7 *Human Mortality Database*. University of California, Berkeley (USAUN), and Max Planck Institute for Demographic Research (Germany). Data downloaded 01/11/2014 from www.mortality.org.

## Acknowledgement

This research was supported by National Institute of Health grant R21AG046760 The final publication is available at Springer via http://dx.doi.org/DOI:10.1007/s10522-016-9669-1.

